# Isolating the effect of beat salience on RAS outcomes

**DOI:** 10.1101/2024.10.15.618556

**Authors:** Kristi M. Von Handorf, Ramkumar Jagadeesan, Jessica A. Grahn

## Abstract

Rhythmic auditory stimulation (RAS) is an intervention for gait-disordered populations that involves synchronizing footsteps to regular auditory cues. Previous research has shown that high-groove music (music that induces the desire to move or dance to it) improves gait relative to low-groove music, but how this effect occurs is unclear. Greater beat salience in high-groove music may improve gait because salient beats are easier to synchronize with. Here, we manipulated beat salience by embedding metronome tones to emphasize beat onsets in both high- and low-groove music. We expected that, if beat salience drives gait improvements to high-groove music, then embedding metronome in low-groove music would elicit similar gait improvements (e.g. increased stride velocity). Here, we quantified gait synchronization in terms of period-matching (overall step rate to the cue pace) and phase-matching (individual step onsets to beat onsets). We tested a sample of healthy younger and older adults, with auditory cues matched to 10% faster than baseline. Low-groove music with embedded metronome, compared to without, elicited better period-matching; there were no differences between metronome conditions in high-groove music. These findings suggest gait improvements to high-groove music could be due to its high beat salience. On the other hand, embedded metronome did not improve phase-matching accuracy, but high-groove music did. This suggests that beat salience may not improve gait via easing step-to-beat synchronization, but rather through an overall increase in movement vigor.

## Introduction

Parkinson’s disease (PD) is a common and debilitating neurodegenerative disorder that affects 1% of adults over age 60 (Samii, Nutt, & Ransom, 2004). In addition to cardinal symptoms (bradykinesia, rigidity, and resting tremor; Postuma, et al., 2015), people with PD exhibit gait impairments such as narrow stride width, unpredictable fluctuations between consecutive strides, and freezing of gait (Ashoori, et al., 2015). Pharmacological treatment and deep-brain stimulation can alleviate cardinal symptoms such as tremor and rigidity, but their benefits on gait impairments are typically limited and decrease over time (Sethi, 2008). Rhythmic auditory stimulation (RAS), which involves walking to regular, isochronous auditory cues, was developed as a relatively cost-free and accessible intervention which has shown promise in improving gait. Typical gait improvements to RAS include longer and faster strides, less variability, and reduced freezing episodes compared to walking in silence (for reviews, see Ghai, et al., 2018; Burrai, et al., 2021; Ye, et al., 2022).

RAS uses music (e.g., Ready, et al., 2022; Roberts, et al., 2021; de Bruin, et al., 2010) or metronome (e.g., Elston, et al., 2010; Howe, et al., 2003; Nieuwboer, et al., 2007). Several studies suggest that “activating” or “motivational” music improves gait outcomes relative to neutral music or metronome-only cues (Styns, et al., 2007; Leman, et al., 2013; de Bruin, et al., 2015; Wittwer, et al., 2013), consistent with movement research in other domains, such as increased time-to-exhaustion when treadmill running (Terry, et al., 2012) and more accurate motor entrainment (Rose, et al., 2019). The idea of “activating” music appears to be related to the property of “groove”, which is defined as the urge to move or dance in response to music (Janata, et al., 2012). In healthy younger and older adults, high-groove music elicits faster and less variable gait than low-groove music (Leow, Parrott, & Grahn, 2014; Ready, et al., 2019; Leow, Rinchon, & Grahn, 2015). Low-groove music can even worsen gait parameters such as stride velocity and length (Ready, et al., 2019).

There is currently no consensus in the literature regarding what features of high-groove (or motivating) music are responsible for gait improvement. Groove is related to a number of musical features, particularly greater enjoyment (Janata, et al., 2012) and increased beat salience, i.e. the clarity of the beat in music (Madison, et al., 2011; Stupacher, et al., 2016). Enjoyment is a plausible explanation for groove’s effects on gait, as music is associated with dopamine release in the reward system (Vuilleumier & Trost, 2015; Salimpoor et al., 2011) and dopamine release is, in turn, associated with increased movement in PD (Mazzoni et al., 2007). However, enjoyment alone does not seem to elicit gait improvement in RAS, at least when the music is unfamiliar (Roberts et al., 2021; Park et al., 2019). Therefore, here we focus on groove’s association with increased beat salience, because higher beat salience has also been associated with more whole-body movement (Burger et al., 2013) as well as more motor synchronization at the beat level (Burger et al., 2014).

The possibility that beat salience may drive groove effects on gait has been underexplored in the literature. Feature extraction studies have shown that beat salience is correlated with higher amounts of movement across various parts of the body, including the feet (Burger, et al., 2013), as well as periodic movement at the beat level (Burger et al., 2014). Additionally, activating music (defined partially by clear marking of the beat) is correlated with faster walking (Leman et al., 2013), possibly because music with a strong beat may be more inviting to move to, which induces quick entrainment to the beat. However, because these studies only measured correlations between beat salience and movement, the musical stimuli varied in multiple dimensions apart from beat salience (e.g. enjoyment); thus it is not clear whether beat salience itself caused changes in movement. A more recent study attempted to isolate the effects of beat salience from other musical confounds by embedding metronome tones to coincide with the beat in selections of high- and low-groove music (Leow, et al., 2021). If beat salience drives the effect of groove on gait, then the effect of groove should be similar when beat salience is equalized across levels of groove. Indeed, Leow et al. found that when a metronome was embedded in the music, stride time and velocity no longer differed between high-groove music and low-groove music. The authors suggested that embedding metronome improved gait because the greater beat salience may reduce the dual-task interference associated with having to synchronize steps to the beat. This dual-task interference would be greater in low-groove than high-groove conditions when no metronome was present, but would be equalized when a metronome was embedded. Therefore, if groove effects are driven by beat salience alone, then, removing the requirement to synchronize should remove this dual-task interference, which should remove gait differences between high and low beat salience as well as high and low groove conditions. Therefore, they also examined gait in trials where participants were not required to synchronize. When free-walking, even though beat salience was equalized, differences between high- and low- groove music remained. These findings suggest that the improvement associated with high-groove music is not solely due to greater ease of synchronization to more salient beats.

However, it may be that beat salience *is* responsible alone for groove improvement, but the measures of synchronization in that study were not sensitive enough to show ease of synchronization. Synchronization is usually quantified by two parameters: the period (matching of step rate to the beat rate of the music cues) and the phase (aligning of step onsets with individual beat onsets). The previous study (Leow, et al., 2021) measured synchronization only via phase-matching, not period-matching. Period-matching was not analysed because music cues were presented at the same pace as participants’ preferred walking rate, so it was not meaningful to assess how well participants were matching step rate to cues that were already matched to their step rate. However, assessing period-matching may be informative because it could reveal gait improvement that phase-matching would not (e.g. high step-to-step variability in matching steps to beat onsets, but overall faster speed relative to baseline). That is, if cues are presented at a different rate than baseline, it’s possible to match the overall rate successfully without matching the phase of each beat, but the reverse is less likely (if steps are consistently in-phase, then the rate is probably also matched). Using a synchronization measure that is easier to achieve is important in the context of gait synchronization, which can be a difficult task. Generally, synchronization does not occur unless explicit instructions to synchronize are given (Leow, et al., 2018; Hove, et al., 2012; Mendonça, et al., 2014; Wittwer, et al., 2013), and phase-matching in the previous study (Leow, et al., 2021) was poor when instructions to synchronize were not given. On the other hand, explicit instructions can worsen gait for populations who are sensitive to dual-task interference, such as older adults (Hausdorff, 2009; Springer et al., 2006) or poor beat-perceivers (Leow, et al., 2014), especially if cued with low-groove music (Leow, et al., 2021; Leow, et al., 2018; Ready, et al., 2019). It’s possible that when beat salience is enhanced with a metronome, people may synchronize even without intention, and potentially avoid the negative effects associated with intentional synchronization.

Thus, in the current study, we used music with a tempo matched to 10% faster than baseline, which allows us to examine the presence of spontaneous or intentional synchronization, and measured both phase- and period-matching. Moreover, previous research has shown that when RAS is cued at 10% faster than baseline, gait speed and variability improves more than when cued at baseline (Hausdorff, 2009; Erra et al., 2019; Cha, Kim, & Chung, 2014). Therefore, we conducted a conceptual replication of Leow, et al.’s (2021) design, but instead cueing participants with music played at 10% faster than baseline pace, either with metronome embedded or without. Synchronization may also be affected by how instructions are given. In the previous study, instructions to synchronize were interleaved (i.e., every trial switched between synchronize and free-walk). Leow et al. (2021) found that instructions to synchronize to the beat elicited slower and more variable strides compared to free-walking, but carryover effects may have masked true differences between instructions. Thus, in the current study, instructions to synchronize were manipulated between-subjects.

To summarize, the main aims of the study were as follows. First, to evaluate the possibility that the higher beat salience in high-groove music drives improvements in gait, we overlaid metronome beats on a selection of high- and low-groove music. If high beat salience is primarily responsible for gait improvements observed in response to high-groove music, then enhancing beat salience (i.e., adding a metronome) to low-groove music should approximate the improvements in gait with high-groove music. Because high-groove music generally also has high beat salience, we expected an interaction between groove and beat salience (embedded metronome), where music with embedded metronome would improve gait more in low-groove music than high-groove music. Here, gait improvements are defined compared to baseline: longer stride length, faster stride velocity, and less variability (measured by coefficient of variation). Second, we aimed to compare the effects of enhanced beat salience when participants were and were not instructed to synchronize to the beat of the music at 10% faster than baseline. We hypothesized that poor beat-perceivers might synchronize poorly in phase-matching but may show at least partial period-matching, especially when the beat was more salient (i.e. embedded metronome). In contrast, we expected good beat-perceivers to both phase-match and period-match successfully, especially when instructed to synchronize.

We tested these hypotheses in both healthy younger and older adults, as older adults may be more sensitive to altered cognitive demands during gait. Gait variability increases with attentional demand during gait in healthy older adults and in PD (Al-Yahya, et al., 2011; Harrison & Earhart, 2023). Thus, examining gait changes in older adults may be more informative for clinical contexts.

## Method

### Participants

Of 100 total participants, 44 young adults from the University of Western Ontario (18-29 years, *M* = 21, *SD* = 3.1, 81% female) and 45 older adults from the London, Ontario community (51-78 years, *M* = 63, *SD* = 7.6, 85% female) were analyzed. Eleven participants were excluded from analyses due to testing error (corrupted or missing data, cadence not altered correctly). No participants reported neurological or hearing disorders. Demographic data by subgroup is presented in Table 1. This study was approved by the Health Sciences Research Ethics Board at the University of Western Ontario. All participants provided informed consent.

**Table 1.** Demographic data by subgroup *(N=89)*. Data presented as means (standard deviations) for age, music training, and dance training. Sums are presented for gender (male/female).

|  | Older Adults |  |  |  | Younger Adults |  |  |  |
| --- | --- | --- | --- | --- | --- | --- | --- | --- |
|  | Free-walking |  | Synchronized |  | Free-walking |  | Synchronized |  |
|  | Good BP | Poor BP | Good BP | Poor BP | Good BP | Poor BP | Good BP | Poor BP |
| N | 13 | 10 | 13 | 9 | 11 | 11 | 12 | 10 |
| Age | 64.5<br>(8.6) | 64.9<br>(6.8) | 60.8<br>(7.1) | 63.7<br>(8.2) | 21.5<br>(3.8) | 20.5<br>(2.6) | 21.6<br>(3.2) | 20.9 (3.0) |
| Gender<br>(M/F) | 1/12 | 1/9 | 4/9 | 0/9 | 1/10 | 4/7 | 1/11 | 1/9 |
| Music<br>Training | 5.2<br>(5.3) | 0.4 (1.0) | 2.2 (4.9) | 0.8 (1.6) | 6.5 (4.3) | 1.3 (2.3) | 3.9 (4.9) | 3.2 (4.2) |
| Dance<br>Training | 2.5<br>(4.3) | 0.1 (0.3) | 0.7 (2.5) | 0.5 (1.3) | 1.0 (1.7) | 3.4 (5.6) | 2.1 (3.0) | 0.6 (1.3) |
| BAT | 78% | 51% | 73% | 54% | 78% | 53% | 76% | 49% |
| Baseline<br>Cadence | 117 | 117 | 110 | 111 | 109 | 106 | 112 | 110 |

### Stimuli

Stimuli were a set of 10 non-lyrical music tracks identical to those used in previous work (Leow, et al., 2021). In the previous study, each music track had a cowbell sound embedded to coincide with the beat, resulting in a total of 20 tracks (10 songs with no metronome embedded, and the same 10 songs with metronome embedded). Metronome-only sequences were also identical to those used in previous work, created using 50-ms 1-kHz sine tones (Leow et al., 2021). All stimuli were played at 10% faster than baseline cadence. Tempo was altered from original versions using the Change Tempo function in Audacity (Audacity® version 2.4.2), which preserves pitch.

Participants rated 15-second clips of each track (both with and without metronome embedded; already altered to 10% faster than baseline) on groove, enjoyment, beat salience, and familiarity. All four ratings were completed before moving on to the next stimulus. The rating scale items were as follows:

1. *Groove:* **How much did the music make you want to move?** 1 = I would definitely not move to this, 50 = I feel indifferent about moving to this, 100 = I would move a lot to this.
2. *Familiarity:* **How familiar are you with this piece of music?** 1 = not at all familiar (never heard it before), 50 = I know this song - I am certain I have heard it before, 100 = I know the song so well that I can predict what happens next in the song.
3. *Enjoyment:* **How much do you enjoy listening to this piece of music?** 1 = I strongly dislike this song, 50 = I feel neutral about this song, 100 = I strongly enjoy this song.
4. *Beat salience:* **How strong is the beat in this piece of music?** 1= very weak, 50 = neutral, 100 = very strong.

Based on individual participants’ ratings, a customized list of songs to walk to was generated for each participant. Of the rated clips, the four songs rated highest on groove and the four songs rated lowest on groove were used for the high- and low-groove conditions, respectively. Each participant walked to 16 songs (4 high-groove, metronome-embedded; 4 high-groove, no metronome; 4 low-groove, metronome-embedded; 4 low-groove, no metronome) and 4 metronome-only tracks.

### Apparatus

Spatiotemporal gait measures were recorded using a 16 ft. Zeno® pressure sensor walkway (Protokinetics, Havertown, PA, USA), at a sampling rate of 120 Hz and spatial resolution of 1.27cm. The walkway was 579 x 90.2cm, with an active sensor area 488 x 61cm. Stimuli were played over wireless headphones (Sennheiser® HDR 160) during both the computer tasks and walking tasks. To ensure precise alignment between auditory stimuli and recorded gait data, a custom-written MATLAB script was triggered to begin playback for each trial when it received a 5V TTL pulse from the Protokinetics walkway.

### Materials

We used the Perception subtest of the Beat Alignment Test (BAT) from the Goldsmiths Musical Sophistication Index v1.0 (Müllensiefen, et al., 2014), which is modelled after the original Beat Alignment Test (Iversen, 2008). The BAT assesses beat perception in the context of music, which is more directly relevant to the current task than other beat perception measures such as the BAASTA (Dalla Bella et al., 2016) and Harvard BAT (Fujii & Schlaug, 2013). In the test, metronome beeps were superimposed over instrumental music clips, and participants determined whether the metronome was correctly aligned with the perceptual beat. Participants completed 3 practice trials and 17 test trials, and stimulus order was randomized for each participant.

### Procedure

To determine baseline cadence (number of steps per minute), each participant walked six lengths of the pressure-sensitive walkway at a comfortable pace, with no auditory stimuli present. To obtain steady-state gait, participants walked to a floor marking 1.78m beyond the edge of the walkway before turning and re-rentering the walkway. Then, participants completed the ratings task, followed by the BAT (using E-Prime 3.0 software; Psychology Software Tools, Pittsburgh, PA) and a demographics questionnaire. Participants walked to a total of 20 tracks (16 music, four metronome-only) on the pressure walkway.

### Data analysis

#### Synchronization performance

Because the music stimuli were from real performances, and thus not perfectly isochronous, canonical beat times were determined using BeatRoot software (Dixon, 2007) for every song at 110 BPM. We used a custom-written MATLAB script to generate a metronome track for each song (both the metronome-embedded and no-metronome version) and then overlaid each metronome track onto its respective song. Trained musicians listened to each track to confirm alignment between music and metronome. On occasion, BeatRoot made small errors in alignment or intended metrical level. When this occurred, the song was entered again into BeatRoot at a different tempo (significantly faster or slower), which, for reasons not clear to the authors, then gave correct canonical times (as verified by trained musicians); all IBIs were then transformed back to 110 BPM. We used the 110 BPM reference grid for all songs, which was then transformed to the corresponding tempo for analysis (i.e. the tempo that participant tapped or walked to, 10% faster than baseline).

In accordance with previous synchronization literature (tapping: Kirschner & Tomasello, 2009; Sowinski & Dalla Bella, 2013; Dalla Bella & Sowinski, 2015, gait: Leow et al., 2017; Dotov et al., 2017; McIntosh et al., 1997), we applied circular statistics to both tapping and gait data. Procedures common to both sets of data are described in this section, and procedures specific to the respective task (tapping or gait) are described in the relevant section below. Using circular measures allows examination of when each response occurs relative to its target beat time set to 0° on the circle. Points on the circle therefore represent the time at which the response occurred relative to the beat (*relative phase angle*), with positive values denoting responses later than the beat time, and negative values denoting responses earlier than the beat time, on a scale of -180° to 180°. Therefore, each response in a single trial was represented as a unit vector at its relative phase angle.

To calculate circular measures, we computed the time difference in milliseconds between each response time and its nearest stimulus beat time (*asynchrony*). To then compute the relative phase difference, the asynchrony was multiplied by the momentary rate (1/momentary inter-response interval), and then multiplied by 360° to represent the phase difference in degrees on the circle. Each trial was then summarized by two values: (1) the circular mean of the relative phase angles (i.e. average difference between beat and response times), and (2) phase coherence (i.e. the length of the resultant mean vector). The mean phase angle is a measure of *synchronization accuracy,* and phase coherence is a normalized measure of *synchronization consistency* on a scale of 0-1. Relatively higher values indicate relatively higher consistency (lower variability). For circular plots, we used the mean phase angle for each participant across trials, where the 0°-360° scale was transformed to -180°-180° to show how responses clustered around the 0° target beat time. For analysis, we used mean absolute phase error (mean of the absolute values of each phase difference in a trial), ignoring whether responses were leading or lagging relative to the beat. This is analogous to using a linear measure of absolute asynchrony (Dalla Bella et al., 2017; Rose et al., 2019).

#### Beat alignment test

Beat perception ability was quantified by the proportion of BAT trials correctly identified as “on” or “off” the beat. Scores were exported from E-Prime, then included in statistical analyses as a continuous variable. For graphing purposes, participants were divided into two groups by the median, in which good beat-perceivers scored at or above the median, and poor beat-perceivers scored below the median (in accordance with previous work: Leow et al., 2021; Roberts et al., 2021).

#### Song ratings

Song ratings were analyzed to ensure (1) that groove ratings differed across high- and low-groove music, (2) that perceived beat salience increased when metronome was embedded, and (3) to check whether groove ratings differed across variables that were not controlled in the current study (enjoyment and familiarity). To perform these checks, we conducted two-way Groove x Beat Salience ANOVAs on each of the four condition ratings (groove, beat salience, enjoyment, familiarity).

### Tapping

To provide an independent measure of whether beat salience increased when metronome was embedded in music, we examined tapping performance across good and poor beat- perceivers. We conducted two Beat Salience (metronome embedded, no metronome embedded) x Beat Perception (good beat-perceiver, poor beat-perceiver) ANOVAs on phase coherence and absolute phase error, in which improvement is measured by higher phase coherence and lower absolute error.

### Gait: period-matching

Gait kinematics were obtained using ProtoKinetics Movement Analysis Software (PKMAS). Footsteps were pre-processed in PKMAS and then exported to R for analysis. During pre-processing, all partial footsteps (entering and exiting the walkway) were excluded. All 20 walks for each participant were then exported to R. All analyses were performed in R 3.5.2 (R Core Team, 2018), using the following statistical packages: *tidyverse* (Wickham, 2017), *rstatix* (Kassambara, 2023), and *emmeans* (Lenth, 2024).

We tested two between-subject factors: age (young, old), instructions to synchronize (synchronize, free-walk), and two within-subject factors: groove (high, low), beat salience (metronome embedded, no metronome embedded). Beat perception scores were included as a covariate. Thus, we ran 2 x 2 x 2 x 2 mixed-measures ANCOVAs on the following eight spatiotemporal gait parameters: stride velocity, stride length, stride time, stride width, double support time, and variability of stride velocity, stride length, and stride time.

Stride length was defined as the anterior-posterior distance from the first contact location of one step to the first contact location of the next ipsilateral step. Stride velocity was defined as stride length divided by stride time (time interval between the first contact time of one step to the first contact time of the next ipsilateral step). Double limb support time was defined as the time between ipsilateral foot-strike to the previous step’s contralateral toe-off.

Gait parameters were analysed in the following families: **gait speed** (stride velocity, and its constituent components: stride length and stride time), **gait variability** (coefficient of variation of stride velocity, length, and time, where the coefficient of variation is the standard deviation of each gait parameter divided by the mean of that gait parameter); and **bias for stability** (stride width and double support time). Bonferroni adjustments were applied to each family. The resulting critical *p*-values were: 0.017 (gait speed), 0.017 (gait variability), and 0.025 (bias for support).

To assess how gait changed relative to baseline, normalized change scores were calculated for each gait parameter, given by the following function:

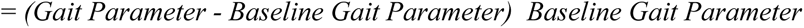

### Gait: phase-matching

Step onsets (first contact time of each step) were analyzed to examine phase-matching. Similar to period-matching analyses, we tested two between-subject factors: age (young, old), instructions to synchronize (synchronize, free-walk), and two within-subject factors: groove (high, low), beat salience (metronome embedded, no metronome embedded). Beat perception scores were included as a covariate. We ran 2 x 2 x 2 x 2 x 2 mixed-measures ANCOVAs on the phase coherence (to measure synchronization consistency) and absolute phase angle error (to measure synchronization accuracy). Bonferroni adjustments were applied to the two ANCOVAs as a family (similar to period-matching measures); thus the resulting critical *p*-value was 0.025 for each test.

## Results

### Song Ratings

Song ratings are shown in Figure 1 across levels of groove and beat salience, and ANOVA results are shown in Table 2. Compared to low-groove songs, high-groove songs were rated significantly higher on groove, beat salience, enjoyment, and familiarity. Songs with an embedded metronome were rated significantly higher on beat salience (as expected), but significantly *lower* on groove and enjoyment. Therefore, any effects of an embedded metronome on gait are likely due to increased beat salience, not increased groove or enjoyment. Additionally, there was an interaction between groove and beat salience for ratings of beat salience, in which an embedded metronome increased beat salience in low-groove music, but not high-groove music. This suggests that embedding a metronome was successful in increasing beat salience, at least on a perceptual level.

**Fig. 1.**
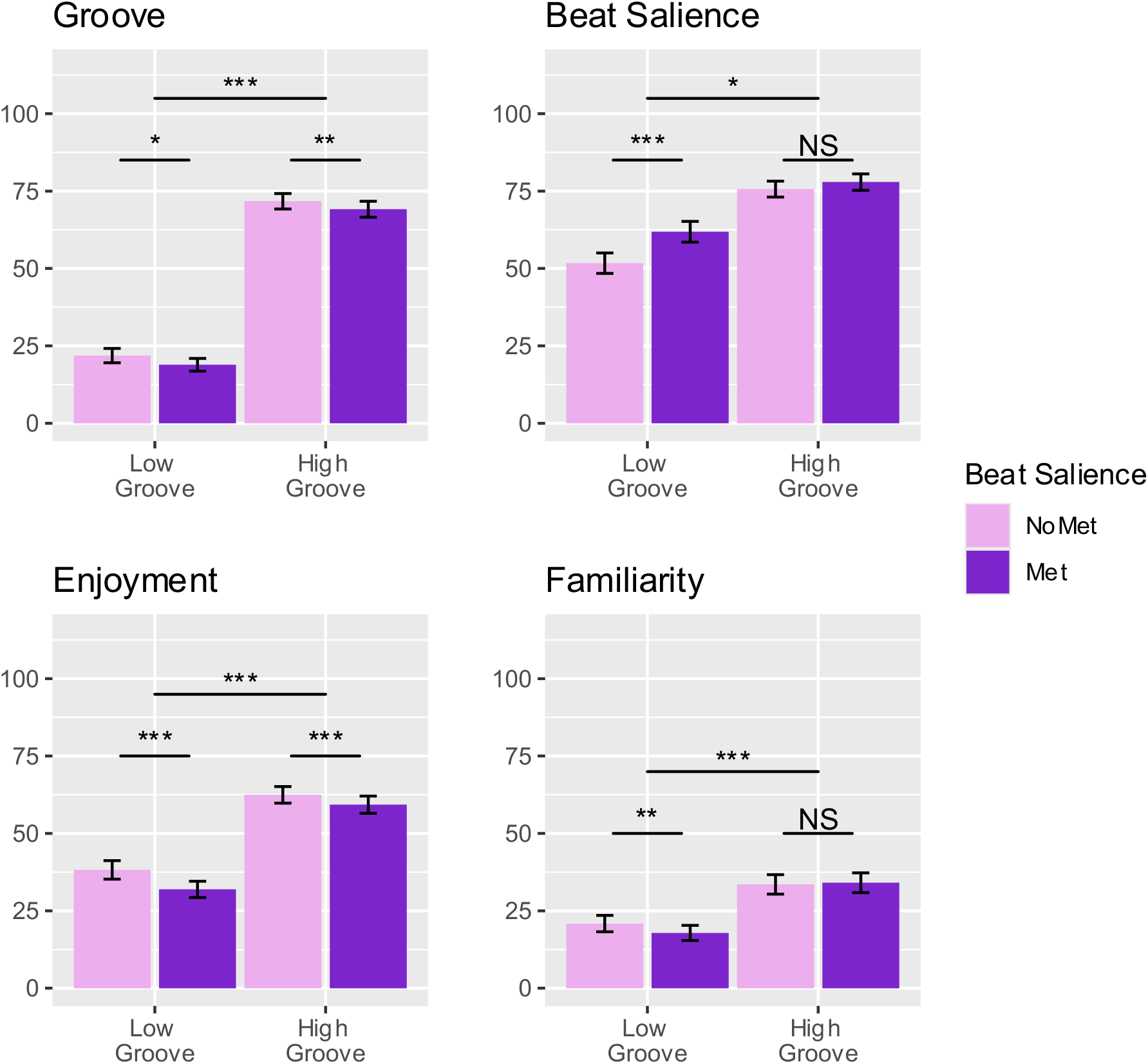
Means of song ratings by condition for music that participants walked to. Music with metronome embedded was rated lower on groove and enjoyment than music without metronome. Embedded metronome in low-groove music significantly increased beat salience ratings, whereas embedded metronome in high-groove music did not.

**Table 2.** ANOVA table of main effects and interactions for each song rating type.

| | *Factor* | *df* | *F* | $\eta^2p$ | *p* |
| --- | --- | --- | --- | --- | --- |
| **Groove** |  |  |  |  |  |
|  | **Groove** | 84 | 586.13 | 0.87 | **<0.001** |
|  | **Beat Salience** | 84 | 10.02 | 0.11 | **0.002** |
|  | Groove X Beat Salience | 84 | 0.42 | 0.00 | 0.52 |
| **Beat Salience** |  |  |  |  |  |
|  | **Groove** | 84 | 100.18 | 0.54 | **<0.001** |
|  | **Beat Salience** | 84 | 16.29 | 0.16 | **<0.001** |
|  | **Groove X Beat Salience** | 84 | 10.39 | 0.11 | **0.002** |
| **Enjoyment** |  |  |  |  |  |
|  | **Groove** | 84 | 141.45 | 0.63 | **<0.001** |
|  | **Beat Salience** | 84 | 15.80 | 0.16 | **<0.001** |
|  | **Groove X Beat Salience** | 84 | 4.30 | 0.05 | **0.04** |
| **Familiarity** |  |  |  |  |  |
|  | **Groove** | 84 | 49.10 | 0.37 | **<0.001** |
|  | Beat Salience | 84 | 1.92 | 0.02 | 0.17 |
|  | Groove X Beat Salience | 84 | 3.31 | 0.04 | 0.07 |

### Tapping

#### Mean absolute phase error

Taps were significantly more aligned to the beat when a metronome was embedded in the music than not, *F*(1, 69) = 5.44, *p* = .02, *η²p* = 0.07; Fig 2A). There was no significant main effect of beat perception ability, *F*(1, 69) = 1.36, *p* = .25, *η²p* = 0.02. There was no interaction between beat salience and beat perception ability, *F*(1, 69) = 0.22, *p* = .64, *η²p* = 0.

**Fig 2.**
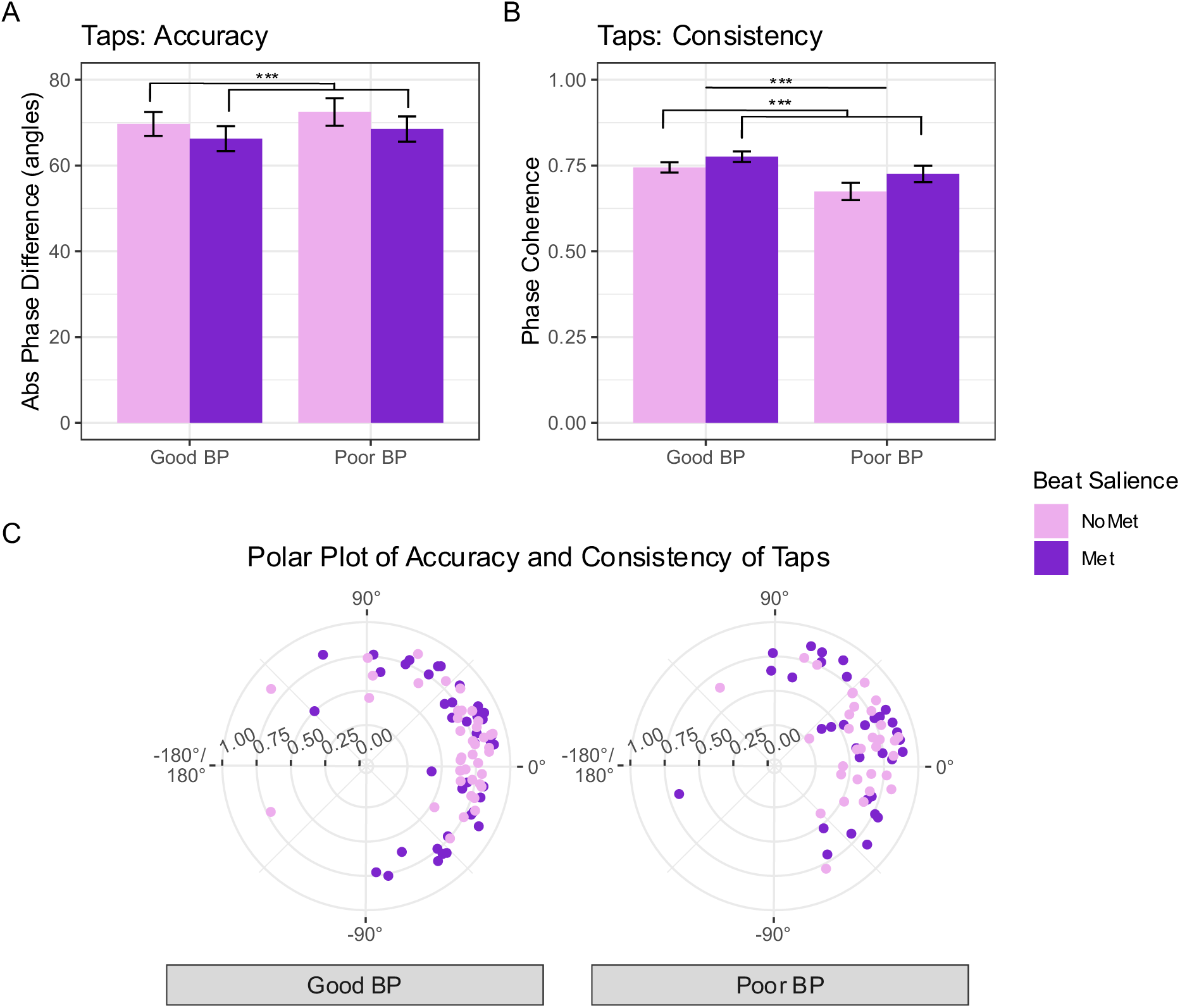
Linear (A, B) and polar (C) plots of accuracy and consistency of taps to beat onsets in music. Plot A shows mean *absolute* phase error per condition (i.e., the data submitted to statistical tests), and Plot B shows the phase coherence between taps and beat onsets in the music. Error bars represent standard error of the mean. Plot C shows both the phase coherence and circular mean of *non-absolute* phase error for each participant (averaged by condition). The mean non-absolute error indicates when taps occurred relative to the beat, where the beat time = 0°; phase angles > 0° are later than the beat; < 0° are earlier than the beat. Points closer to the edge of the circle indicate higher consistency (less variability) in tap times. Music with an embedded metronome increased tapping accuracy and consistency compared to without, in both good and poor beat-perceivers.

### Phase coherence

As expected, music with embedded metronome significantly increased phase coherence, *F*(1, 69) = 26.29, *p* < .001, *η²p* = 0.28; Fig 2B. Good beat-perceivers tapped significantly more consistently than poor beat-perceivers, *F*(1, 69) = 9.02, *p* = .003, *η²p* = 0.12. There was no significant interaction between beat salience and beat perception ability, *F*(1, 69) = 0.03, *p* = .87, *η²p* = 0.

### Gait: Period-Matching Gait Speed

#### Stride velocity

Participants walked significantly faster to high-groove than to low-groove music, *F*(1,78) = 20.18, *p* < .001, *η²p* = 0.2; Fig 3A. They walked significantly faster to music with metronome than without metronome, *F*(1,78) = 11.71, *p* < .001, *η²p* = 0.12. There were no main effects of instructions, age, or beat perception ability, and no significant interactions (Table 3).

**Fig. 3.**
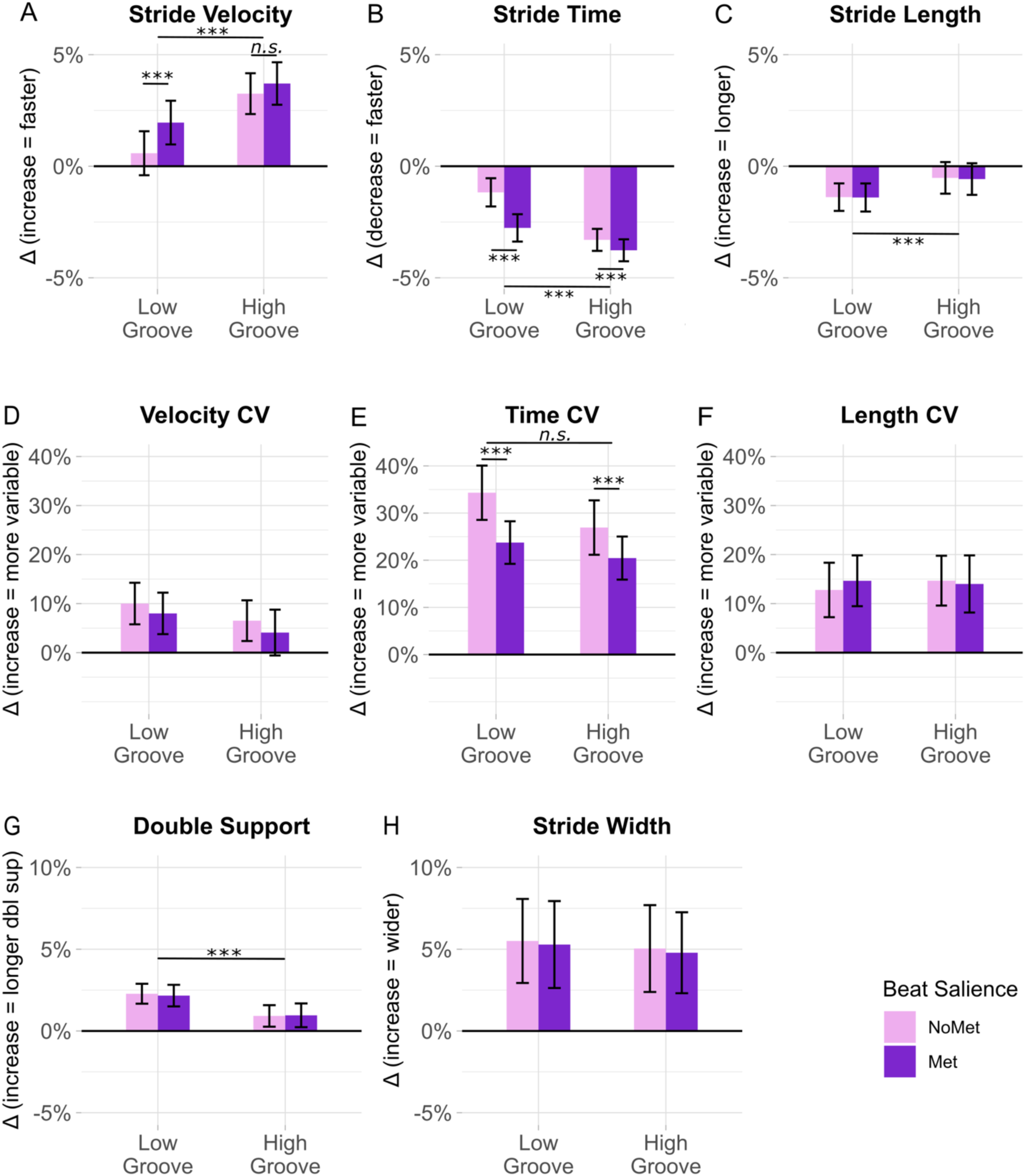
Effects of groove and beat salience on gait, collapsed across instructions, beat perception ability, and age. All gait parameters (e.g. stride velocity) are plotted relative to the baseline (percent change). Error bars represent standard error of the mean. \*\**p* < .01, \*\*\**p* < .001, and n.s. = not significant. High-groove music elicited faster and longer strides, and less bias for support, than low-groove music. Music with metronome elicited faster and less variable strides compared to music without metronome, especially in low-groove music.

**Fig. 4.**
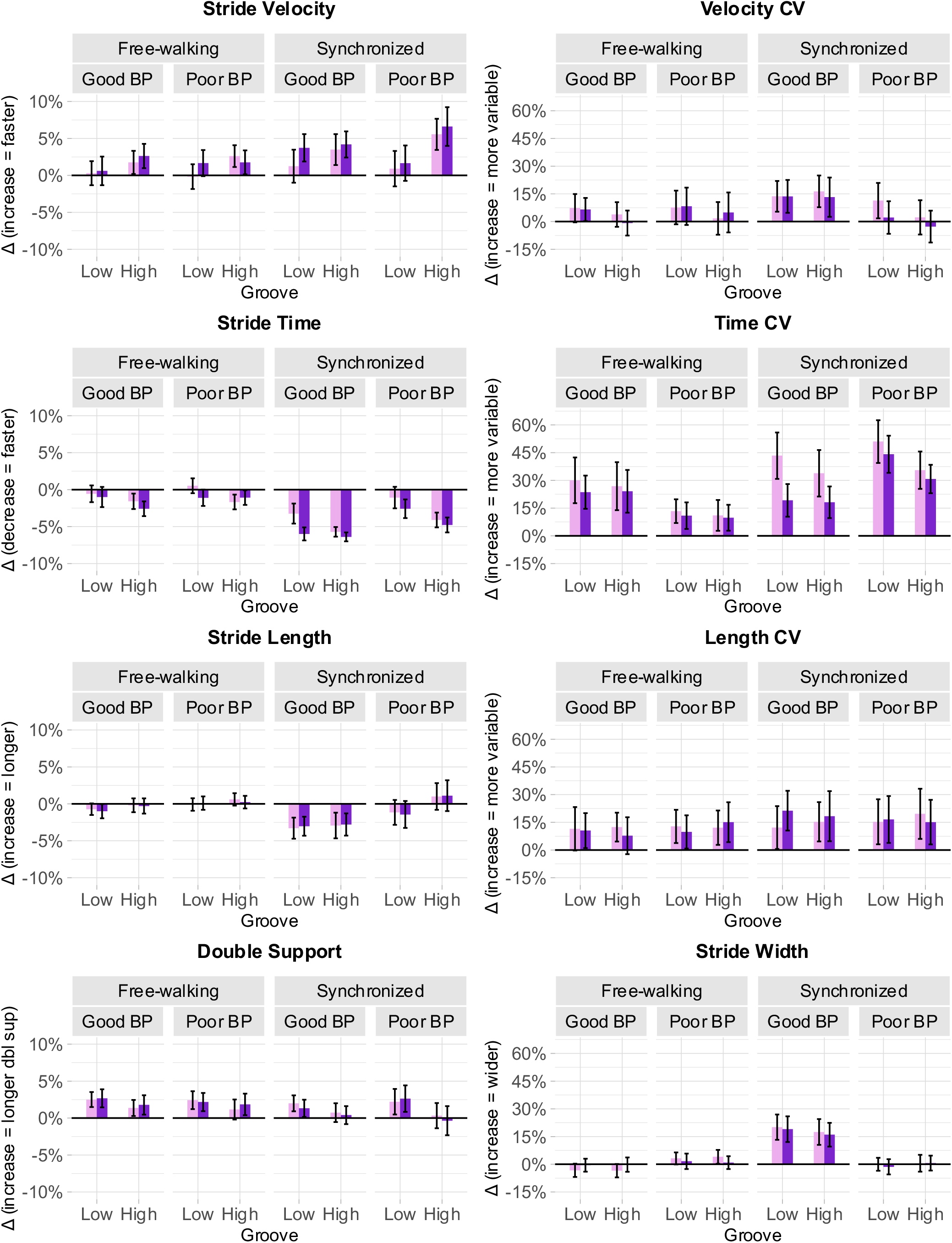
Effects of groove and beat salience on gait, split across instructions to synchronize and beat perception ability levels. All gait parameters (e.g. stride velocity) are plotted relative to the baseline (percent change). Error bars represent standard error of the mean. Participants walked significantly faster when instructed to synchronize compared to free-walk. Good and poor beat-perceivers were split at the median for visualization, but beat perception scores were included as a continuous variable for analysis. There were no significant effects of beat perception ability.

**Fig. 5.**
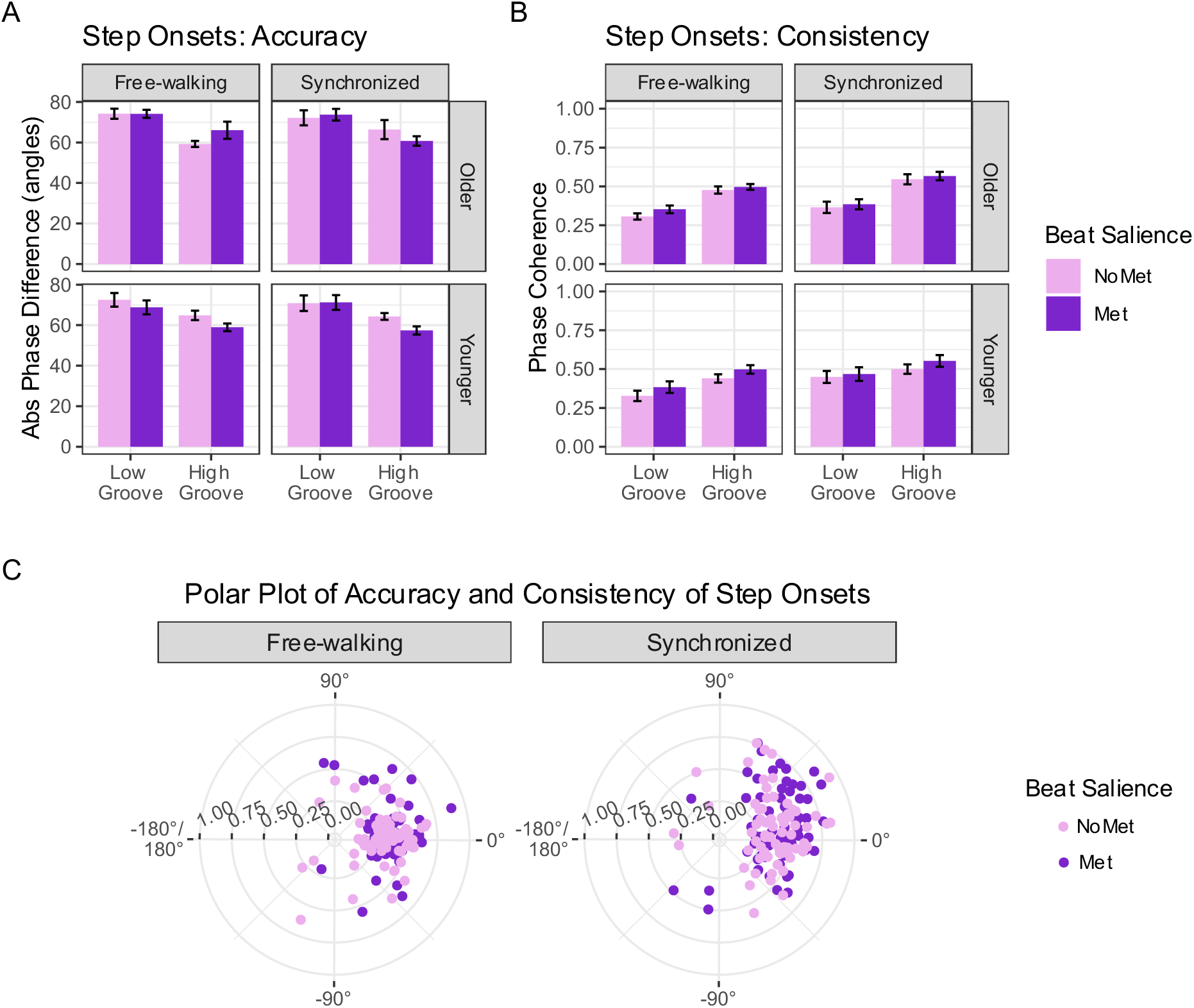
**Linear (A, B) and polar (C) plots of phase-matching accuracy and consistency of step onsets to beat onsets in music.** Plot A shows mean *absolute* phase error per condition (i.e., the data submitted to statistical tests), and Plot B shows the phase coherence between step onsets and beat onsets in the music. Error bars represent standard error of the mean. Plot C shows both the phase coherence and circular mean of *non-absolute* phase error for each participant (averaged by condition). The mean non-absolute error indicates when step onsets occurred relative to the beat, where the beat time = 0°; phase angles > 0° are later than the beat; < 0° are earlier than the beat. Points closer to the edge of the circle indicate higher consistency (less variability) in step onset times. High-groove music increased accuracy, but higher beat salience (embedded metronome) and instructions to synchronize did not. Higher beat salience, high-groove music, and instructions to synchronize increased consistency, but there were no interactions between these factors, suggesting that beat salience improved consistency of phase-matching separately from its impact on groove.

**Table 3.**
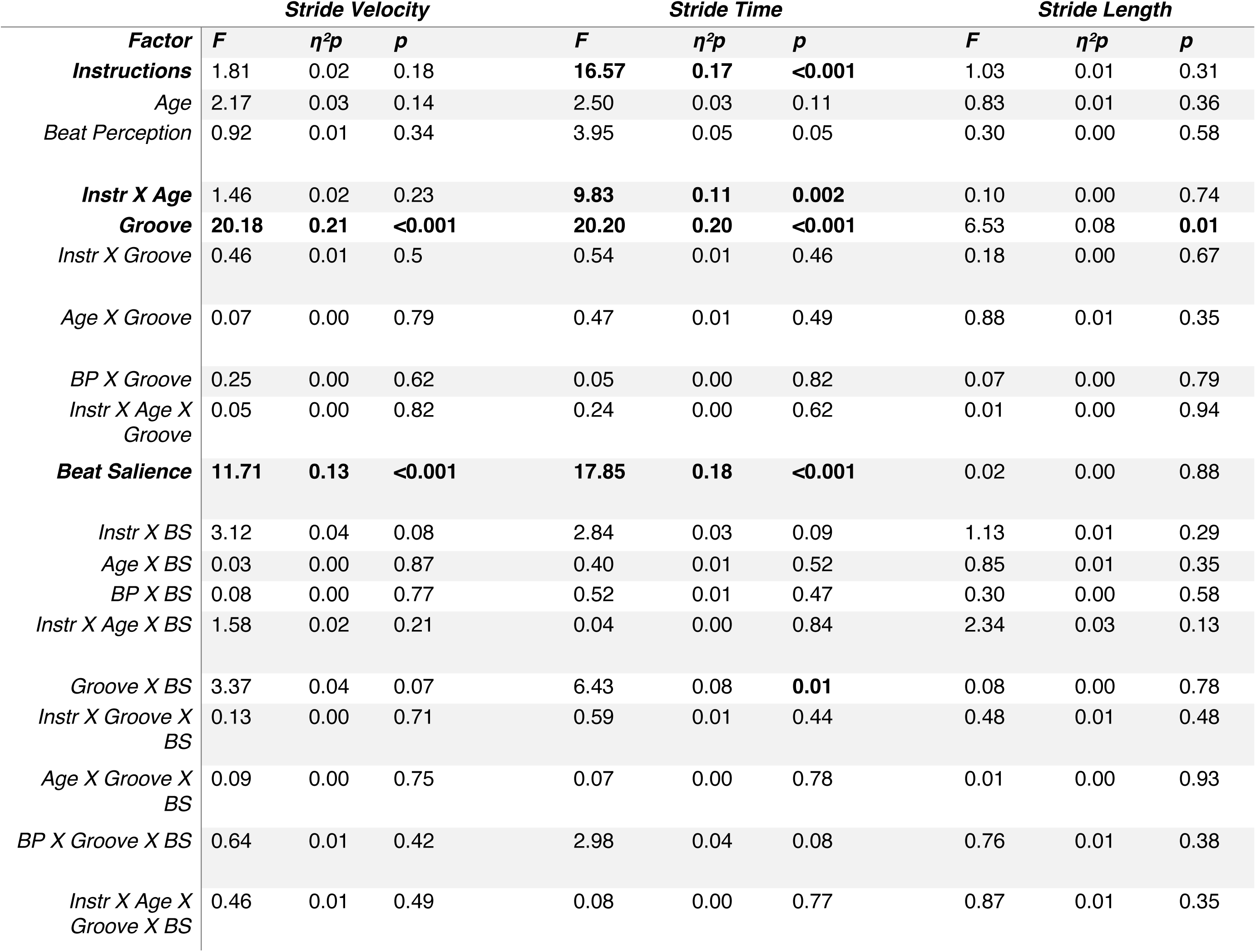
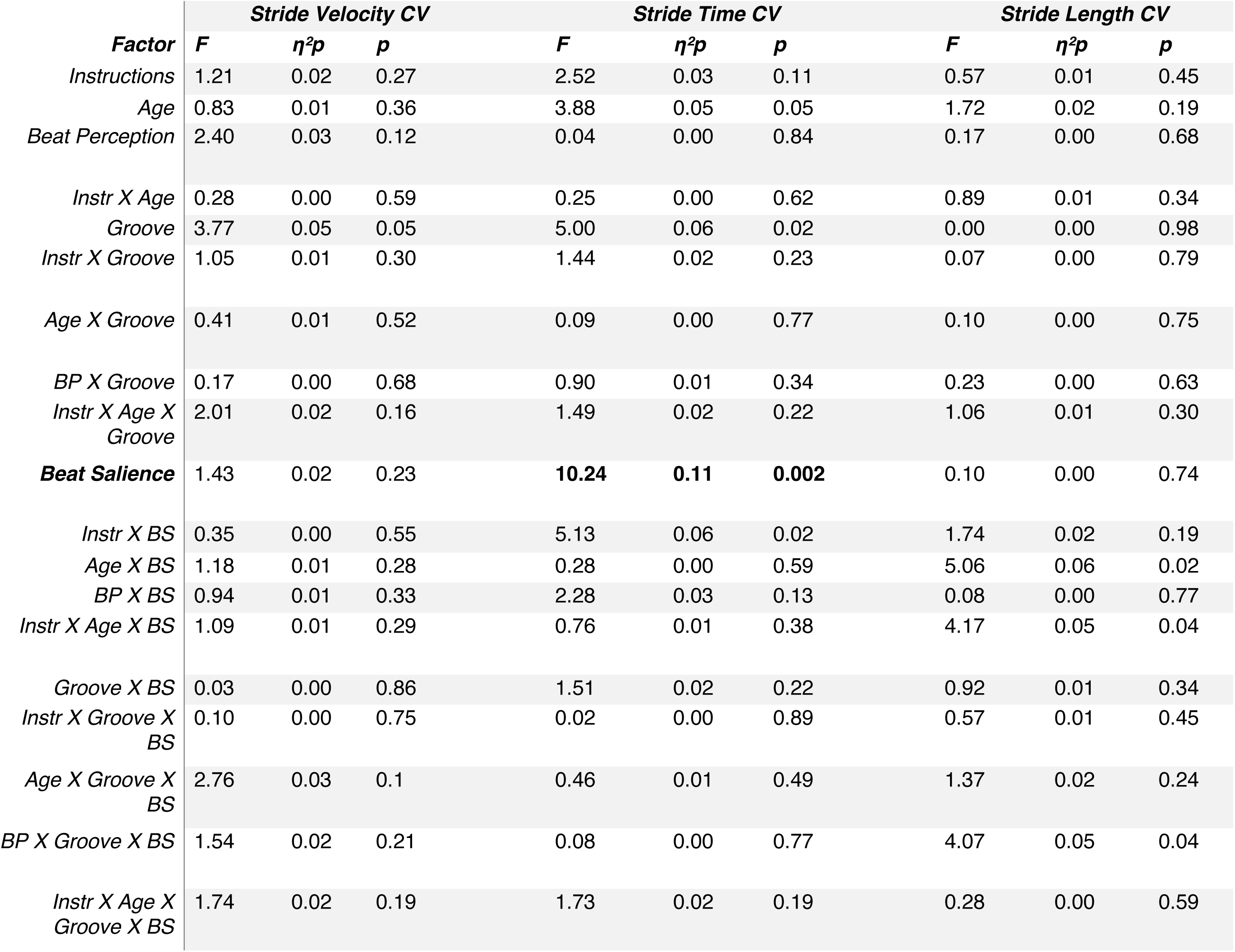
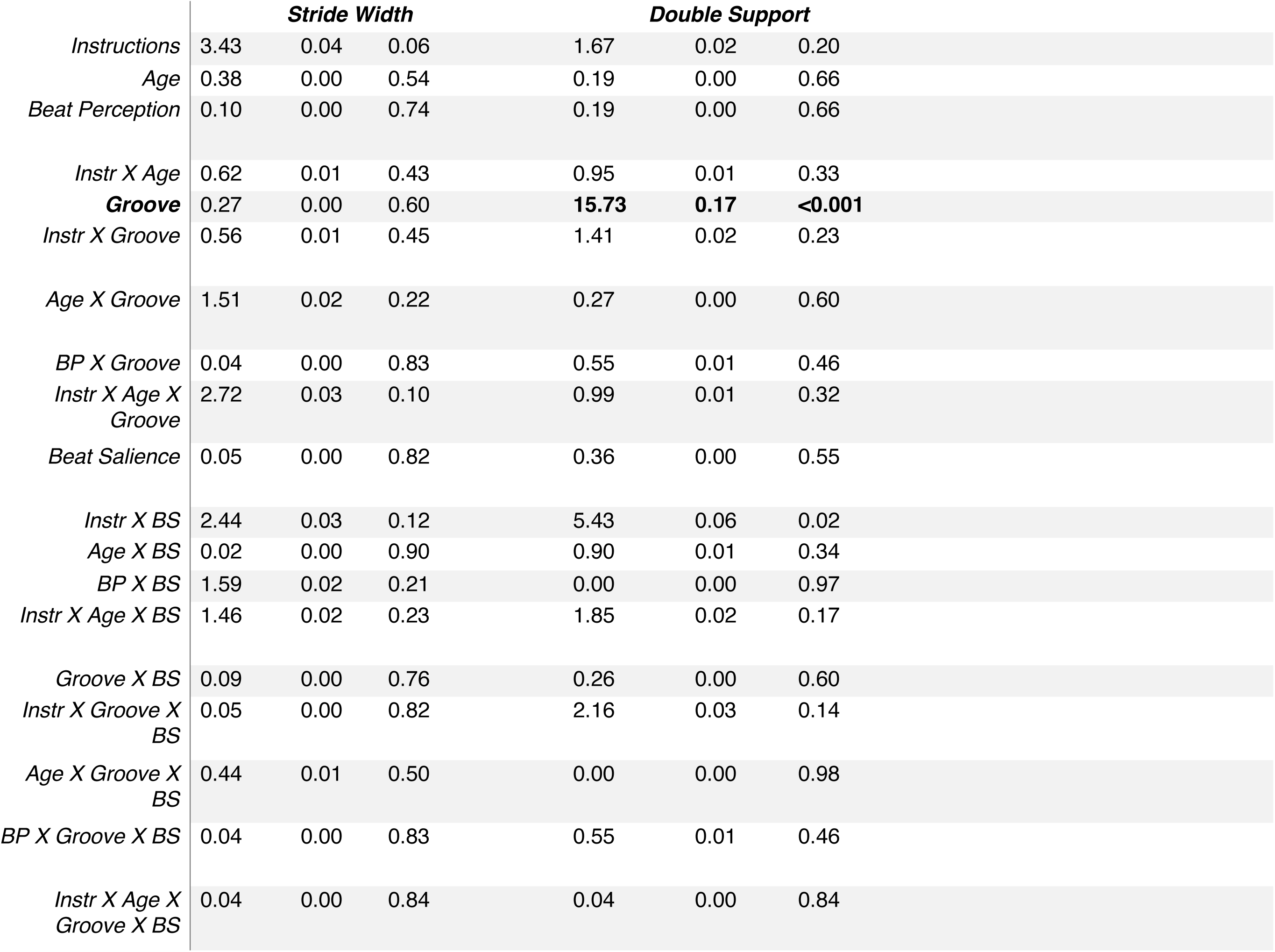
ANOVA table of main effects and interactions for each gait parameter.

### Stride time

Participants had a significantly faster step rate to high-groove than to low-groove music, *F*(1,78) = 20.20, *p* < .001, *η²p* = 0.20; Fig 3B. They also had a significantly faster step rate to music with metronome than without metronome, *F*(1,78) = 17.85, *p* < .001, *η²p* = 0.18). There was a significant interaction between groove and beat salience, *F*(1,78) = 6.43, *p* = .01, *η²p* = 0.08, in which music with metronome elicited a faster step rate than music without metronome in *both* high-groove and low-groove music, but the difference in stride time was greater in low-groove music.

Instructions to synchronize elicited a faster step rate than free-walking, *F*(1,78) = 16.57, *p* < .001, *η²p* = 0.17. This was qualified by an interaction between instructions and age, *F*(1,78) = 9.83, *p* = .002, *η²p* = 0.1, where older adults walked faster when synchronizing than free-walking, but younger adults did not differ between instructions. There were no significant main effects of beat perception ability and no other significant interactions (Table 3).

### Stride length

Participants took significantly longer strides to high-groove than to low-groove music, *F*(1,78) = 6.53, *p* = .01, *η²p* = 0.08; Fig 3C. There were no main effects of beat salience, instructions, age, or beat perception ability, and no significant interactions (Table 3).

### Gait variability

There were no significant effects of groove, beat salience, age, beat production ability, or instructions on **stride velocity variability** (Fig 3D; Table 3) or **stride length variability** (Fig 3F; Table 3). However, there was a significant effect of beat salience on **stride time variability**. Participants walked with significantly lower stride time variability to music with metronome than music without metronome, *F*(1,78) = 10.24, *p* = .002, *η²p* = 0.11; Fig 3E. Also, the effect of groove on stride time variability approached significance, *F*(1,78) = 5.0, *p* = .02, *η²p* = 0.06. There were no significant effects of age, instructions to synchronize, or beat perception ability (Table 3).

### Bias for support

There were no significant effects of groove, beat salience, age, instructions to synchronize, or beat perception ability on **stride width** (Fig 3H; Table 3). There was a significant main effect of groove on **double-limb support time,** where DLST was significantly greater for low-groove music than high-groove music, *F*(1,78) = 15.73, *p* < .001, *η²p* = 0.17; Fig 3G. There were no other main effects or interactions for DLST (Table 3).

### Gait: Phase-Matching

#### Mean absolute phase error

Step onsets were significantly more aligned to the beat in high-groove than low-groove music, *F*(1, 58) = 23.42, *p* < .001, *η²p* = 0.29; Fig 5A. However, there was no significant effect of beat salience, *F*(1, 58) = 3.3, *p* = .07, *η²p* = 0.05. Instructions to synchronize did not improve accuracy of step onsets, *F*(1, 58) = 0.05, *p* = .81, *η²p* = 0, nor did beat perception ability, *F*(1, 58) = 1.32, *p* = .25, *η²p* = 0.02. An interaction between groove, beat salience, and instructions neared significance, *F*(1, 58) = 4.89, *p* = .03, *η²p* = 0.07.

### Phase coherence

Phase coherence was significantly greater for high-groove than low-groove music, *F*(1, 58) = 80.79, *p* < .001, *η²p* = 0.58; Fig 5B. Phase coherence was also significantly greater to music with metronome than no metronome, *F*(1,58) = 9.77, *p* = .003, *η²p* = 0.14, and when instructed to synchronize vs. free-walk, *F*(1, 58) = 8.52, *p* = .005, *η²p* = 0.13). There were no significant main effects of beat perception ability, *F*(1, 58) = 0.12, *p* = .73, *η²p* = 0, or age, *F*(1, 58) = 0.16, *p* = .69, *η²p* = 0.

There was an interaction between age and groove, *F*(1, 58) = 7.49, *p* < .008, *η²p* = 0.11, where older adults were more negatively affected by low-groove music (low-groove: *M* = 0.35, *SE* = 0.02; high-groove: *M* = 0.53, *SE* = 0.02) than younger adults (low-groove: *M* = 0.41, *SE* = 0.02; high-groove: *M* = 0.5, *SE* = 0.02). There were no other significant interactions.

## Discussion

### Effect of increasing beat salience in low- and high-groove music

We examined whether the higher beat salience that accompanies high-groove music is responsible for improvements in gait (Madison et al., 2011; Stupacher et al., 2016). We expected an interaction between groove and beat salience, where low-groove music with metronome would improve gait compared to without metronome, but high-groove music would not differ (due to inherently high beat salience). Indeed, we found this interaction in stride time, in which music with metronome elicited a faster step rate than music without metronome, especially in low-groove music. Additionally, stride velocity increased and stride time variability decreased overall when metronome was embedded (regardless of groove). These findings were similar to a previous study by Leow et al. (2021) in which participants walked to music that was set to their baseline pace (compared to 10% faster than baseline in the current study). In that study, adding a metronome to low-groove music increased stride velocity compared to without metronome, and there were no differences for high-groove music with and without metronome. In the current study, this interaction was found in stride time; this difference may be because, in their attempt to synchronize their footsteps to the accelerated cue rate (10% faster), participants mainly altered their stride time. Regardless, the fact that gait velocity increased when metronome was added to low-groove music is consistent with the idea that high beat salience is an important contributor to gait improvements to high-groove music.

In addition to assessing gait synchronization overall, we also measured phase-matching performance. We found that step-to-beat synchronization accuracy was better for high-groove music compared to low-groove music, but beat salience (embedded metronome) did *not* significantly affect accuracy. On the other hand, synchronization consistency increased for both high-groove and metronome-embedded music, although there was no interaction. These results together suggest that although increased beat salience improves consistency, the effect of groove is much larger, and beat salience makes very little difference for accuracy. Therefore, it appears that high-groove music improves synchronization at both the period and phase, but beat salience is not responsible for improvement at the phase, at least not alone. Previous literature on groove perception suggests that beat salience is not the only feature of rhythm that may facilitate gait; for example, moderate levels of syncopation (rhythmic events that violate listener expectations) often lead to higher groove ratings (Witek et al., 2014; Matthews et al., 2022; Spiech et al., 2022). Thus, simply highlighting every beat with a metronome (as in the current study) may not be enough to facilitate synchronization to the beat, compared to a pattern of emphasis that also takes into account the metrical level (a simple example would be to emphasize the onsets of only the second and fourth beats in a four-beat measure).

Interestingly, although embedded metronome did not improve phase-matching, it *did* improve gait overall relative to baseline (faster strides, with less variability, compared to no metronome). This difference between phase-matching and period-matching performance suggests that embedded metronome may improve gait through a mechanism other than facilitating synchronization of steps to beat onsets. Beat salience has been proposed to improve gait via an increase in overall movement vigor (Leow et al., 2021; Park et al., 2019), although it seems unlikely that this movement increase occurs because of greater reward, because enjoyment ratings went down with embedded metronome in the current study.

### Effect of beat perception and instructions to synchronize

The second aim was to compare effects of increased beat salience when participants were and were not instructed to synchronize to the beat of the music. Because of increased dual-task sensitivity, we hypothesized that poor beat-perceivers might show poor phase-matching when asked to synchronize, but that period-matching may be easier, especially when the beat is more salient (i.e. embedded metronome). However, in contrast to previous literature (Ready et al., 2022; Roberts et al., 2021; Ready et al., 2019; Leow et al., 2014), we found no differential effects of beat salience for good or poor beat perceivers, and this was true both when participants free-walked or synchronized to the music. In fact, we found no significant difference in any gait parameters between good and poor beat-perceivers. Previous work (Leow et al., 2021) found that poor beat- perceivers took more variable strides when instructed to synchronize, especially in low-groove music. The negative effect of synchronizing to music may have been overcome because poor beat- perceivers were instructed to walk 10% faster than their natural pace.

Overall, synchronizers tended to take faster and similarly variable strides relative to free- walkers. These results are consistent with previous findings that synchronizing to music improves temporal gait parameters in healthy adults (Ready et al., 2022; Leow et al., 2015; Mendonça et al., 2014). However, several studies have found that synchronization elicits slower and shorter strides compared to baseline (Roberts et al., 2021; Leow et al., 2018; Ready et al., 2019), or that overall variability increases even if gait speed improves (Ready et al., 2022). This may be because synchronizing gait is cognitively demanding, especially because it requires whole body coordination to synchronize (compared to finger tapping). We speculate that synchronization improved gait in the current study because participants were cued at 10% faster than normal walking pace (versus 15% or 22.5% faster in other studies), because 10% faster allows them to achieve faster velocity while avoiding the difficulty of synchronizing to a rate that is too far from their normal pace.

### Cue pace

The goal of cueing participants at 10% faster than baseline was to elicit faster gait velocity. Here, participants tended to increase gait speed relative to baseline, regardless of instructions to synchronize, in contrast to previous work that presented cues at baseline pace, in which participants took slower and shorter strides relative to baseline (Leow et al., 2021). Our findings are consistent with that of other studies with music-based RAS with cue rates different from preferred stepping rate (Ready et al., 2022; Leow et al., 2014; Leow et al., 2015; Roberts et al., 2021; Erra et al., 2019; Cha, Kim, & Chung, 2014), suggesting that temporal gait characteristics may improve in response to the speed of the music, even if the full speed is not achieved (10% in this case).

### Clinical implications

#### High-groove music as optimal stimulus

We replicated previous findings that high-groove music elicits faster and longer strides than low-groove music (Leow et al., 2021; Ready et al., 2019; Leow et al., 2014; Styns et al., 2007; Leman et al., 2013; de Bruin et al., 2015). Metronome-only cues elicited similar gait improvements as high-groove music, which is also consistent with previous research (Ready et al., 2022; de Bruin et al., 2015). Here, strides still tended to shorten relative to baseline, but stride length was maintained better overall in the current study than when cued at baseline pace (Leow et al., 2021), possibly due to the faster cueing pace. Moreover, high-groove music elicited smaller increases in velocity variability and double support than low-groove music. These effects were found regardless of age, beat perception ability, or instructions to synchronize. This study adds to the body of literature that suggests high-groove music may be optimal in RAS therapy. Although metronome-only cues and high-groove music have similar effects on gait, there may be a motivational benefit to listening to music over metronome cues, potentially facilitating long-term adherence.

### Individual differences in beat perception

Variable responses to RAS cues have been linked to individual differences in rhythmic ability among patient populations (Dalla Bella et al., 2017; Dalla Bella et al., 2018; Cochen de Cock et al., 2018). Specifically, better beat perception is associated with higher likelihood of positive response to RAS, possibly because it eases synchronization to the beat, although participants in these studies were not explicitly instructed to synchronize (Dalla Bella et al., 2018). Here, we found that poor beat-perceivers did not alter gait when free-walking, and tended to walk faster overall when synchronizing. This is in contrast to other suggestions that poor beat-perceivers may take slower and shorter strides because they experience an attentional cost associated with synchronizing to music (Leow et al., 2021; Roberts et al., 2021). Future studies are needed before strong clinical recommendations regarding rhythmic abilities can be made.

## References

Ashoori, A., Eagleman, D. M., & Jankovic, J. (2015). Effects of auditory rhythm and music on gait disturbances in parkinson’s disease. Frontiers in Neurology, 6. 10.3389/fneur.2015.00234

Audacity Team (2014). Audacity(R): Free Audio Editor and Recorder [Computer program]. Version 2.4.2 retrieved September 26th 2020 from http://audacity.sourceforge.net/

Dalla Bella, S., Benoit, C.-E., Farrugia, N., Keller, P. E., Obrig, H., Mainka, S., & Kotz, S. A. (2017). Gait improvement via rhythmic stimulation in Parkinson’s disease is linked to rhythmic skills. Scientific Reports, 7(1). 10.1038/srep42005

Dalla Bella, S., Dotov, D., Bardy, B., & de Cock, V. C. (2018). Individualization of music-based rhythmic auditory cueing in Parkinson’s disease. Annals of the New York Academy of Sciences, 1423(1), 308–317. 10.1111/nyas.13859

Burger, B., Thompson, M. R., Luck, G., Saarikallio, S., & Toiviainen, P. (2013). Influences of rhythm- and timbre-related musical features on characteristics of music-induced movement. Frontiers in Psychology, 4. 10.3389/fpsyg.2013.00183

Burrai, F., Apuzzo, L., & Zanotti, R. (2021). Effectiveness of rhythmic auditory stimulation on gait in parkinson disease. *Holistic Nursing Practice*, Publish Ahead of Print. 10.1097/hnp.0000000000000462

Cha, Y., Kim, Y., & Chung, Y. (2014). Immediate effects of rhythmic auditory stimulation with tempo changes on gait in stroke patients. Journal of Physical Therapy Science, 26(4), 479–482. 10.1589/jpts.26.479

Cochen De Cock, V., Dotov, D., Geny, C., Bardy, B., Driss, V., & Dalla Bella, S. (2018). Rhythmic abilities and musical training in Parkinson’s disease: Do they help? Annals of Physical and Rehabilitation Medicine, 61, e101. 10.1016/j.rehab.2018.05.216

Dalla Bella, S. (2015). Using music to improve mobility in Parkinson’s disease: Effects beyond gait? Annals of Physical and Rehabilitation Medicine, 58, e71.

Dalla Bella, S., Farrugia, N., Benoit, C.-E., Begel, V., Verga, L., Harding, E., & Kotz, S. A. (2016). BAASTA: Battery for the assessment of auditory sensorimotor and timing abilities. Behavior Research Methods, 49(3), 1128–1145. 10.3758/s13428-016-0773-6

Dalla Bella, S., & Sowiński, J. (2015). Uncovering beat deafness: Detecting rhythm disorders with synchronized finger tapping and perceptual timing tasks. Journal of Visualized Experiments, 97. 10.3791/51761-v

de Bruin, N., Doan, J. B., Turnbull, G., Suchowersky, O., Bonfield, S., Hu, B., & Brown, L. A. (2010). Walking with music is a safe and viable tool for gait training in parkinson’s disease: The effect of a 13-week feasibility study on single and dual task walking. Parkinson’s Disease, 2010, 1–9. 10.4061/2010/483530

de Bruin, N., Kempster, C., Doucette, A., Doan, J. B., Hu, B., & Brown, L. A. (2015). The effects of music salience on the gait performance of young adults. Journal of Music Therapy, 52(3), 394–419. 10.1093/jmt/thv009

Dotov, D. G., Bayard, S., Cochen de Cock, V., Geny, C., Driss, V., Garrigue, G., Bardy, B., & Dalla Bella, S. (2017). Biologically-variable rhythmic auditory cues are superior to isochronous cues in fostering natural gait variability in Parkinson’s disease. Gait & Posture, 51, 64–69. 10.1016/j.gaitpost.2016.09.020

Elston, J., Honan, W., Powell, R., Gormley, J., & Stein, K. (2010). Do metronomes improve the quality of life in people with Parkinson’s disease? A pragmatic, single-blind, randomized cross-over trial. Clinical Rehabilitation, 24(6), 523–532. 10.1177/0269215509360646

Erra, C., Mileti, I., Germanotta, M., Petracca, M., Imbimbo, I., De Biase, A., Rossi, S., Ricciardi, D., Pacilli, A., Di Sipio, E., Palermo, E., Bentivoglio, A. R., & Padua, L. (2019). Immediate effects of rhythmic auditory stimulation on gait kinematics in Parkinson’s disease ON/OFF medication. Clinical Neurophysiology, 130(10), 1789–1797. 10.1016/j.clinph.2019.07.013

Etani, T., Marui, A., Kawase, S., & Keller, P. E. (2018). Optimal Tempo for Groove: Its Relation to Directions of Body Movement and Japanese nori. Frontiers in Psychology, 9. 10.3389/fpsyg.2018.00462

Fujii, S., & Schlaug, G. (2013). Harvard beat assessment test. PsycTESTS Dataset. 10.1037/t36474-000

Ghai, S., Ghai, I., Schmitz, G., & Effenberg, A. O. (2018). Effect of rhythmic auditory cueing on parkinsonian gait: A systematic review and meta-analysis. Scientific Reports, 8(1). 10.1038/s41598-017-16232-5

Kassambara, A. (2023). rstatix: Pipe-Friendly Framework for Basic Statistical Tests. R package version 0.7.2, <https://CRAN.R-project.org/package=rstatix>.

Harrison, E. C., & Earhart, G. M. (2023). The effect of auditory cues on gait variability in people with parkinson’s disease and older adults: A systematic review. Neurodegenerative Disease Management, 13(2), 113–128. 10.2217/nmt-2021-0050

Hausdorff, J. M. (2009). Gait dynamics in Parkinson’s disease: Common and distinct behavior among stride length, gait variability, and fractal-like scaling. Chaos: An Interdisciplinary Journal of Nonlinear Science, 19(2). 10.1063/1.3147408

Hove, M. J., Suzuki, K., Uchitomi, H., Orimo, S., & Miyake, Y. (2012). Interactive rhythmic auditory stimulation reinstates natural 1/f timing in gait of parkinson’s patients. PLoS ONE, 7(3), e32600. 10.1371/journal.pone.0032600

Howe, T. E., Lövgreen, B., Cody, F. W., Ashton, V. J., & Oldham, J. A. (2003). Auditory cues can modify the gait of persons with early-stage Parkinson’s disease: A method for enhancing parkinsonian walking performance? Clinical Rehabilitation, 17(4), 363–367. 10.1191/0269215503cr621oa

Iversen, J. (2008). Neural dynamics of beat perception and production. The Journal of the Acoustical Society of America, 124(4_Supplement), 2431–2431. 10.1121/1.4808956

Janata, P., Tomic, S. T., & Haberman, J. M. (2012). Sensorimotor coupling in music and the psychology of the groove. Journal of Experimental Psychology: General, 141(1), 54–75. 10.1037/a0024208

Kirschner, S., & Tomasello, M. (2009). Joint drumming: Social context facilitates synchronization in preschool children. Journal of Experimental Child Psychology, 102(3), 299–314. 10.1016/j.jecp.2008.07.005

Leman, M., Moelants, D., Varewyck, M., Styns, F., van Noorden, L., & Martens, J.-P. (2013). Activating and relaxing music entrains the speed of beat synchronized walking. PLoS ONE, 8(7), e67932. 10.1371/journal.pone.0067932

Lenth R (2024). emmeans: Estimated Marginal Means, aka Least-Squares Means. R package version 1.10.2, <https://CRAN.R-project.org/package=emmeans>.

Leow, L., Rinchon, C., & Grahn, J. (2015). Familiarity with music increases walking speed in rhythmic auditory cuing. Annals of the New York Academy of Sciences, 1337(1), 53–61. 10.1111/nyas.12658

Leow, L.-A., Parrott, T., & Grahn, J. A. (2014). Individual differences in beat perception affect gait responses to low- and high-groove music. Frontiers in Human Neuroscience, 8. 10.3389/fnhum.2014.00811

Leow, L.-A., Waclawik, K., & Grahn, J. A. (2017). The role of attention and intention in synchronization to music: Effects on gait. Experimental Brain Research, 236(1), 99–115. 10.1007/s00221-017-5110-5

Leow, L.-A., Watson, S., Prete, D., Waclawik, K., & Grahn, J. A. (2021). How groove in music affects gait. Experimental Brain Research, 239(8), 2419–2433. 10.1007/s00221-021-06083-y

Madison, G., Gouyon, F., Ullén, F., & Hörnström, K. (2011). Modeling the tendency for music to induce movement in humans: First correlations with low-level audio descriptors across music genres. Journal of Experimental Psychology: Human Perception and Performance, 37(5), 1578–1594. 10.1037/a0024323

Matthews, T. E., Witek, M. A. G., Thibodeau, J. L. N., Vuust, P., & Penhune, V. B. (2022). Perceived Motor Synchrony With the Beat is More Strongly Related to Groove Than Measured Synchrony. Music Perception, 39(5), 423–442. 10.1525/mp.2022.39.5.423

Mazzoni, P., Hristova, A., & Krakauer, J. W. (2007). Why don’t we move faster? Parkinson’s disease, movement vigor, and implicit motivation. The Journal of Neuroscience, 27(27), 7105–7116. 10.1523/jneurosci.0264-07.2007

Mullensiefen, D., Gingras, B., Musil, J., & Stewart, L. (2014). Goldsmiths musical sophistication index. PsycTESTS Dataset. 10.1037/t42817-000

Nieuwboer, A., Kwakkel, G., Rochester, L., Jones, D., van Wegen, E., Willems, A. M., Chavret, F., Hetherington, V., Baker, K., & Lim, I. (2007). Cueing training in the home improves gait-related mobility in Parkinson’s disease: The RESCUE trial. *Journal of Neurology*, Neurosurgery & Psychiatry, 78(2), 134–140. 10.1136/jnnp.200x.097923

Park, K. S., Hass, C. J., Fawver, B., Lee, H., & Janelle, C. M. (2019). Emotional states influence forward gait during music listening based on familiarity with music selections. Human Movement Science, 66, 53–62. 10.1016/j.humov.2019.03.004

Postuma, R. B., Berg, D., Stern, M., Poewe, W., Olanow, C. W., Oertel, W., Obeso, J., Marek, K., Litvan, I., Lang, A. E., Halliday, G., Goetz, C. G., Gasser, T., Dubois, B., Chan, P., Bloem, B. R., Adler, C. H., & Deuschl, G. (2015). MDS clinical diagnostic criteria for Parkinson’s disease. Movement Disorders, 30(12), 1591–1601. 10.1002/mds.26424

Ready, E. A., Holmes, J. D., & Grahn, J. A. (2022). Gait in younger and older adults during rhythmic auditory stimulation is influenced by groove, familiarity, beat perception, and synchronization demands. Human Movement Science, 84, 102972. 10.1016/j.humov.2022.102972

Ready, E. A., McGarry, L. M., Rinchon, C., Holmes, J. D., & Grahn, J. A. (2019). Beat perception ability and instructions to synchronize influence gait when walking to music-based auditory cues. Gait & Posture, 68, 555–561. 10.1016/j.gaitpost.2018.12.038

Roberts, B. S., Ready, E. A., & Grahn, J. A. (2021). Musical enjoyment does not enhance walking speed in healthy adults during music-based auditory cueing. Gait & Posture, 89, 132–138. 10.1016/j.gaitpost.2021.04.008

Rose, D., Delevoye-Turrell, Y., Ott, L., Annett, L. E., & Lovatt, P. J. (2019). Music and metronomes differentially impact motor timing in people with and without parkinson’s disease: Effects of slow, medium, and fast tempi on entrainment and synchronization performances in finger tapping, toe tapping, and stepping on the spot tasks. Parkinson’s Disease, 2019, 1–18. 10.1155/2019/6530838

Salimpoor, V. N., Benovoy, M., Larcher, K., Dagher, A., & Zatorre, R. J. (2011). Anatomically distinct dopamine release during anticipation and experience of peak emotion to music. Nature Neuroscience, 14(2), 257–262. 10.1038/nn.2726

Samii, A., Nutt, J. G., & Ransom, B. R. (2004). Parkinson’s disease. The Lancet, 363(9423), 1783–1793. 10.1016/s0140-6736(04)16305-8

Sethi, K. (2008). Levodopa unresponsive symptoms in Parkinson disease. Movement Disorders, 23(S3), S521–S533. 10.1002/mds.22049

Sowiński, J., & Dalla Bella, S. (2013). Poor synchronization to the beat may result from deficient auditory-motor mapping. Neuropsychologia, 51(10), 1952–1963. 10.1016/j.neuropsychologia.2013.06.027

Spiech, C., Sioros, G., Endestad, T., Danielsen, A., & Laeng, B. (2022). Pupil drift rate indexes groove ratings. Scientific Reports, 12(1). 10.1038/s41598-022-15763-w

Stupacher, J., Hove, M. J., & Janata, P. (2016). Audio features underlying perceived groove and sensorimotor synchronization in music. Music Perception, 33(5), 571–589. 10.1525/mp.2016.33.5.571

Styns, F., van Noorden, L., Moelants, D., & Leman, M. (2007). Walking on music. Human Movement Science, 26(5), 769–785. 10.1016/j.humov.2007.07.007

Terry, P. C., Karageorghis, C. I., Saha, A. M., & D’Auria, S. (2012). Effects of synchronous music on treadmill running among elite triathletes. Journal of Science and Medicine in Sport, 15(1), 52–57. 10.1016/j.jsams.2011.06.003

Vuilleumier, P., & Trost, W. (2015). Music and emotions: From enchantment to entrainment. Annals of the New York Academy of Sciences, 1337(1), 212–222. 10.1111/nyas.12676

Wickham, H., Averick, M., Bryan, J., Chang, W., McGowan, L.D., François, R., Grolemund, G., Hayes, A., Henry, L., Hester, J., Kuhn, M., Pedersen, T.L., Miller, E., Bache, S.M., Müller, K., Ooms, J., Robinson, D., Seidel, D.P., Spinu, V., Takahashi, K., Vaughan, D., Wilke, C., Woo, K., & Yutani, H. (2019). “Welcome to the tidyverse.” Journal of Open Source Software, 4(43), 1686. <10.21105/joss.01686>.

Witek, M. A. G., Clarke, E. F., Wallentin, M., Kringelbach, M. L., & Vuust, P. (2014). Syncopation, body-movement and pleasure in groove music. PLoS ONE, 9(4), e94446. 10.1371/journal.pone.0094446

Wittwer, J. E., Webster, K. E., & Hill, K. (2013). Music and metronome cues produce different effects on gait spatiotemporal measures but not gait variability in healthy older adults. Gait & Posture, 37(2), 219–222. 10.1016/j.gaitpost.2012.07.006

Ye, X., Li, L., He, R., Jia, Y., & Poon, W. (2022). Rhythmic auditory stimulation promotes gait recovery in Parkinson’s patients: A systematic review and meta-analysis. Frontiers in Neurology, 13. 10.3389/fneur.2022.940419

